# Maturation of GABAergic signalling times the opening of a critical period in *Drosophila melanogaster*

**DOI:** 10.1101/2025.04.24.650400

**Authors:** Jack Corke, Mariam Huertas Radi, Matthias Landgraf, Richard A. Baines

## Abstract

The occurrence of critical periods during the development of neural networks is widely documented. Activity manipulation when these periods are open can lead to permanent, and often debilitating, effects to the mature neural network. Detailed understanding of the specific contribution of critical periods to network development, however, remains elusive. This is partly because identified critical periods in mammals are present in complex sensory networks (e.g., visual and auditory) that make focused experimental manipulation challenging. It is significant, therefore, that critical periods have been identified in simpler model systems. A critical period occurs during the development of the embryonic locomotor network in the fruit fly, *Drosophila melanogaster*. Perturbation of neuronal activity during this period is sufficient to permanently destabilise the mature larval locomotor network: leaving it prone to induced seizures. Given a clear role of γ-aminobutyric acid (GABA) in the timing of the mammalian critical period of ocular dominance, we sought to establish whether this neurotransmitter also regulates the opening of the *Drosophila* locomotor critical period. Utilising GABA agonists, antagonists, and genetics, we manipulated the embryonic GABAergic system and, at the end of larval life, measured an induced seizure phenotype in mature third-instar larvae. We found that potentiating GABAergic signalling, via embryonic exposure to diazepam or overexpression of the GABA_A_ receptor *rdl*, induced precocious opening of the critical period. By contrast, exposure to the GABA antagonist gabazine, or knockdown of the GABA-synthetic enzyme *Gad1*, delayed opening. Thus, we show that critical period timing within the *Drosophila* CNS is dictated by GABAergic signalling, indicating a phylogenetically conserved role.

## Introduction

Developing neural networks exhibit windows of heightened plasticity termed critical periods (CPs) (Hensch, 2005). Activity manipulation during a CP can have profound and permanent consequences for functionality of a mature network (Coulson et al., 2022, Giachello and Baines, 2015, Hensch, 2005). CPs have thus far been identified across a wide range of developing networks, including mammalian visual and auditory cortex, vocalisation in songbirds, and motor systems in rodents (Hensch, 2005, Hensch and Quinlan, 2018, Hubel and Wiesel, 1970, Harrison et al., 2005, Soiza-Reilly and Azcurra, 2009). Despite their almost ubiquitous occurrence, the contribution of a CP to network development and, moreover, how activity perturbation during these periods can encode permanent change to network function, remains poorly understood. A prime reason for this is the complexity of the circuits in which CPs have been identified.

Work by several groups has identified a CP during the development of the larval locomotor network of *Drosophila* (Crisp et al., 2008, Crisp et al., 2011, Fushiki et al., 2013, Giachello and Baines, 2015). The numerical simplicity of the *Drosophila* larval CNS, coupled with the published larval connectome and genetic tools to manipulate identified synaptic pairings (Valdes-Aleman et al., 2021, Giachello et al., 2022, Winding et al., 2023), provides unique experimental tractability. Previous work from our group has shown that activity-manipulation of the embryonic locomotor CP, which occurs between 17-19 h after egg laying (h AEL, 80 - 90% full embryonic development), induces permanent effects to network stability. On the one hand, activity manipulation during the CP, in wildtype embryos, leads to a marked reduction in network stability promoting a ‘seizure-like’ phenotype in response to strong electrical stimulation. By contrast, activity manipulation in a seizure mutant (*para*^*Bss*^) is sufficient to permanently rescue the seizure phenotype regardless of the presence of the mutation (Giachello and Baines, 2015). Follow-on studies show that competing activity levels are integrated during this CP to allow the developing locomotor network to compensate for unpredictable influences and thus to encode an appropriate excitation:inhibition balance (Hunter et al., 2024). Such unpredictable influences include variations in external parameters, (e.g. seasonal temperature differences), as well as mutations.

Before the *Drosophila* locomotor CP can be fully exploited as a favourable model to better understand mammalian CP function, it is essential that the factors that regulate CP opening and closure are identified. Considerable evidence exists to suggest that in mammals maturation of GABAergic inhibitory signalling triggers the opening of CPs. For example, overexpression of BDNF in mice, in parvalbumin (PV)-expressing fast-spiking GABAergic interneurons, advanced their maturation and as part of that increased GAD65 synthesis, thus potentiating GABAergic signalling. This manipulation induced premature opening of the CP for ocular dominance (Hanover et al., 1999). Similarly, diazepam, a benzodiazepine and positive GABA_A_ receptor allosteric modulator (Hosie and Sattelle, 1996), similarly induced precocious opening of the same CP (Fagiolini and Hensch, 2000). Lastly, mice deficient in GAD65, with reduced levels of available GABA, exhibited delayed CP opening. This effect was rescued by exposure to diazepam, highlighting a clear role for GABA in CP opening (Fagiolini and Hensch, 2000).

Here, we report that temporal manipulation of GABAergic signalling, using both pharmacology and genetics, regulates the timing of the *Drosophila* locomotor CP, consistent with opening occurring with maturation of GABAergic signalling. Our results thus further validate the use of this tractable *Drosophila* CP to better understand the role of CPs in development.

## Materials and Methods

### *Drosophila* husbandry and stocks

Flies were maintained in an LD incubator at 25*°*C (12 h light:dark cycle), except for embryos transiently exposed to 32*°*C, after which they were returned to the original light-dark cycle. Flies were fed on a standard cornmeal medium. *Drosophila* strains used in this study are listed in Table 1. Adults were crossed at a ratio of 1:2 of Gal4-line females to UAS-line males.

**Table 1.**
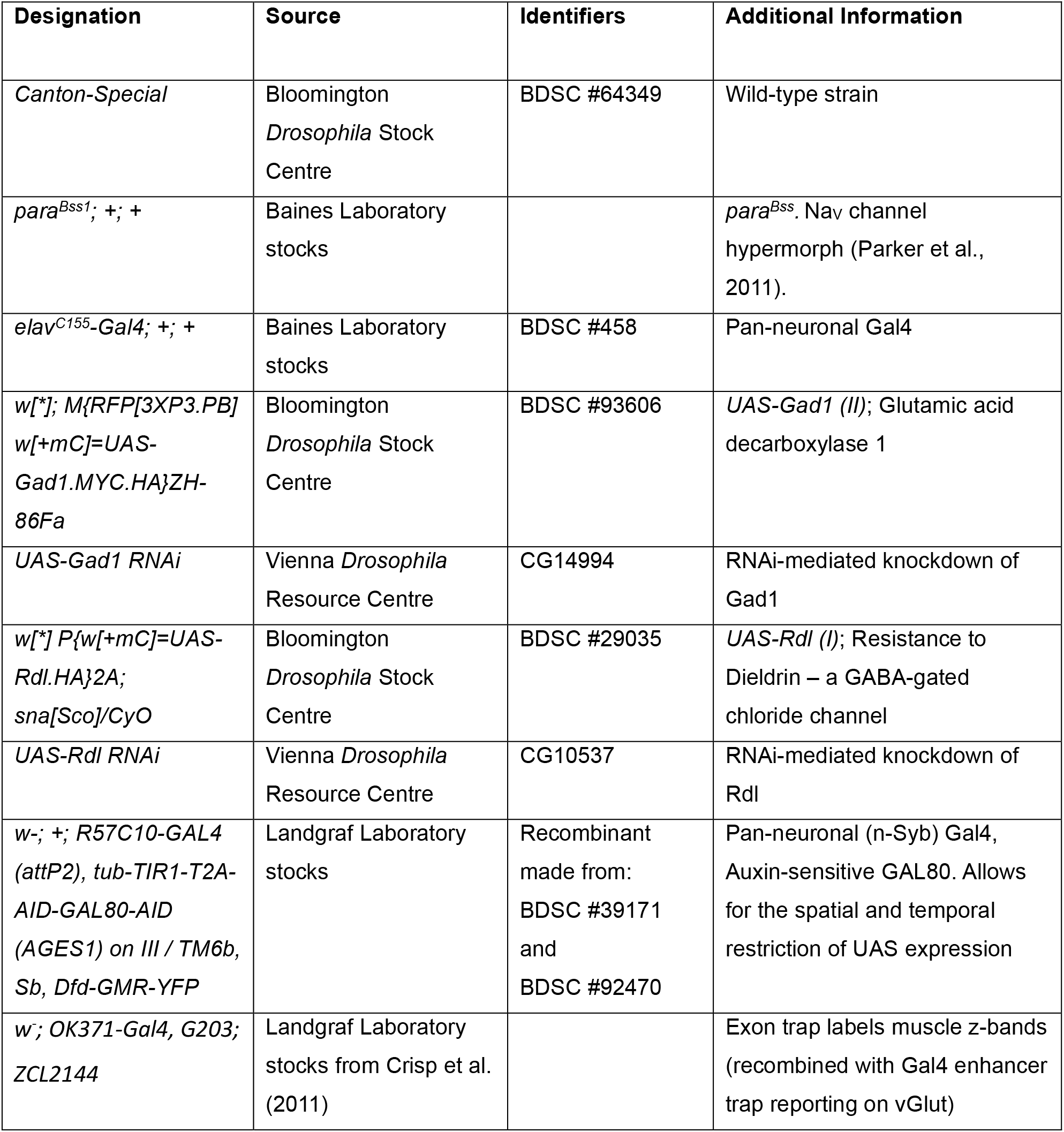
Fly strains used in this study.

### Auxin-inducible, Gal4-compatible gene expression system

The Gal4/UAS expression system ensures spatial control of transgene expression. However, this system does not offer reliable temporal control. The auxin-inducible Gal4-compatible gene expression system (AGES) exploits auxin (potassium 1-naphthaleneacetate) dependent breakdown of a ubiquitously expressed Gal80 repressor that inhibits Gal4. Presence of auxin leads to degradation of the Gal80 repressor, thus allowing Gal4 expression (McClure et al., 2022). Auxin (5 mM, TCI, Japan) was fed to gravid females by mixing with dried yeast paste (Melford, UK) made in water (1 mL ddH_2_O mixed with 4 g yeast extract powder), and allowing flies to feed *ad libitum* for 3 days prior to embryo collection. This protocol has been demonstrated to effect transfer of drug to embryo, which is then metabolised and excreted during early larval development (Marley and Baines, 2011).

### Drug feeding protocol

Drugs were made as stock solutions: diazepam (Sigma-Aldrich, UK) 50 mM stock (in ethanol); gabazine (SR-95561; Sigma-Aldrich, UK) 27 mM (ddH_2_O); potassium 1-naphthaleneacetate (auxin; TCI, Japan) 500 mM stock (ddH_2_O). Final concentrations (0.5 mM diazepam and gabazine, 5 mM Auxin) were made by adding stock to 1 mL ddH_2_O mixed with 4 g yeast extract powder (Melford, UK). 1 mL of this paste was then added to the top of a grape agar plate (containing 6 mL grape agar, Dutscher, UK). Controls were vehicle alone. Mated flies (of both sexes) were allowed to feed on these plates for 3 days to effect transfer of drug to embryo (as described (Marley and Baines, 2011)). Adult flies were contained in 100 mL (15 mm diameter) collection pots kept at 25°C for 3 days to promote egg laying. Plates and food were changed twice daily.

### Embryonic temperature manipulation

Embryos were collected for 1 h time periods and collections were staggered to ensure three separately aged groups could be manipulated simultaneously at 15-17, 17-19 and 19-21 h AEL (± 30 min), respectively. Collected embryos (both sexes) were transferred to 0.2 mL Eppendorf tubes and placed in a PCR machine (Eppendorf Mastercycler Gradient) programmed to maintain the ambient temperature at 25*°*C for 13 h (17:00-06:00), 32*°*C for 2 h (06:00-08:00), then held at 25*°*C thereafter. Manipulated embryos/hatched first instar larvae were transferred into standard cornmeal food vials and reared at 25*°*C until wall-climbing third instar (L3).

### Larval electroshock assay

Wall-climbing L3 larvae (both sexes) were transferred to a clear plastic dish, gently washed in ddH_2_O and dried by paper towel. To administer an electroshock, a stimulator composed of two conductive tungsten wires (0.1 mm diameter), spaced 1-2 mm apart, was gently applied to the larval cuticle (anterior dorsal surface) over the approximate location of the larval CNS. Seizure-like activity was triggered by the application of direct current (6 v, 2 s), provided by an Isolated Voltage Stimulator (DS2A mkII, Digitimer Ltd, UK). Seizure-like activity was defined by immobility, a trembling head, tonic contraction of body wall muscles and the extension of posterior segments (roughly A3 to 9). Seizure recovery was indicated by forward peristaltic waves, resulting in forward crawling. Duration of seizure, i.e., recovery time (RT), was recorded, and reflected the period between stimulus onset and cessation of seizure-like activity. Recordings were limited to 300 s and larvae that failed to recover within this time were discarded from analyses. Videos of this behaviour are available in (Marley and Baines, 2011).

### Embryo staging

Embryos were collected from overnight laying pots. Grape agar plates with embryos were gently rinsed using ddH_2_O, and embryos were transferred into a cell strainer (Falcon 40 µm, Scientific Laboratory Supplies, UK). Cell strainers were then immersed in 50% bleach (sodium hypochlorite in ddH_2_O; Fischer Scientific, UK) for 2 min with gentle shaking, followed by a ddH_2_O rinse. Finally, embryos were transferred to an empty plastic dish with saline (135 mM NaCl [Fisher Scientific, UK], 5 mM KCl [Fisher Scientific], 4 mM MgCl_2_·6H_2_O [Sigma-Aldrich], 2 mM CaCl_2_·2H_2_O [Fisher Scientific], 5 mM TES [Sigma-Aldrich], 36 mM sucrose [Fisher Scientific], at pH 7.15). Early stage 16 embryos (∼13 h AEL) were selected under a microscope (Leica MZ6) upon observation of a ‘three-part gut’, where the gut is divided into three distinct bands. Embryos were kept at 25*°*C in saline until the required developmental age (16, 17, 18 and 19 h AEL) for imaging.

### Imaging peripheral muscle activity

Imaging was performed on w-; OK371-Gal4, ZCL2144 embryos from developmental ages 16 to 19 h AEL. Individual embryos were placed, dorsal side up, in a droplet of saline, on a Sylgard (World Precision Instruments, UK) coated coverslip mounted on a glass microscope slide. The slide was positioned under a compound microscope (BX51WI, Olympus), viewed using a 20x / 0.5 NA water immersion lens (UMPlanFL N, Olympus) and imaged (EXi Aqua camera, QImaging). Blue light (470 nm OptoLED, Cairn Research) was used to visualise GFP, and images recorded for 180 s at 2 fps using Winfluor v4.1.5 (University of Strathclyde). During imaging, embryos were maintained at 25*°*C via a heated plate (CO 102 Digital Heating Unit, Linkam Scientific, UK). Visual analysis of recordings was performed to quantify patterned muscle activity. The subsequent contraction and relaxation of two adjacent z-discs, throughout the length of the embryo, was measured as patterned muscle activity. The mean number of contractions was calculated from 7 to 10 individual embryos per developmental age and treatment group.

### Statistical analysis

Data were analysed using GraphPad Prism v9 (GraphPad Software, San Diego, CA). Data are expressed as mean ± SEM. Sample sizes (N) are reported in figure legends. Full descriptive statistics are reported in Table 2. Shapiro-Wilk tests for normal distribution were conducted on all datasets to determine whether a parametric or non-parametric test was employed. Outliers were identified (ROUT; Q = 1%) and removed from analyses. Statistical tests used included a one-way ANOVA with Tukey’s HSD *post hoc* (parametric) or a Kruskal-Wallis test with Dunn’s *post hoc* (non-parametric) analysis. Significance was defined at *p* ≤ 0.05. The experimenter was blinded to all genotypes and manipulations during both data collection and analysis.

**Table 2.** Full results and statistics.

| Line<br>(Experiment) | Temperature<br>Exposure Age<br>(h AEL) | Mean<br>recovery<br>time (s) ±<br>SEM | Median<br>recovery<br>time (s)<br>(Interquartile<br>Range) | Normally<br>Distributed<br>(Y/N) | post hoc<br>test | p | Figure |
| --- | --- | --- | --- | --- | --- | --- | --- |
| *Canton-S*<br>(CP Timeline) | 15-17 | 65 ± 5 | 61 (25) | Y | Kruskal-Wallis,<br>Dunn's | 0.0001<br>(****) | 1 |
|  | 17-19 | 167 ± 14 | 140 (61) | N |  |  |  |
|  | 19-21 | 59 ± 5 | 57 (18) | Y |  |  |  |
|  | 25°C | 76 ± 6 | 81 (43) | Y |  |  |  |
| *Canton-S*<br>(diazepam) | 15-17 | 154 ± 14 | 138 (123) | N | Kruskal-Wallis,<br>Dunn's | 0.0002<br>(***) | 2A |
|  | 17-19 | 80 ± 5 | 84 (39) | Y |  |  |  |
|  | 19-21 | 99 ± 12 | 79 (87) | Y |  |  |  |
|  | 25°C | 97 ± 8 | 86 (50) | N |  |  |  |
| *Canton-S*<br>(diazepam<br>control) | 15-17 | 82 ± 5 | 83 (30) | N | Kruskal-Wallis,<br>Dunn's | 0.0001<br>(****) | 2B |
|  | 17-19 | 171 ± 13 | 166 (47) | Y |  |  |  |
|  | 19-21 | 80 ± 5 | 81 (40) | Y |  |  |  |
|  | 25°C | 81 ± 5 | 78 (60) | N |  |  |  |
| *Canton-S*<br>(gabazine) | 15-17 | 84 ± 5 | 83 (31) | Y | Kruskal-Wallis,<br>Dunn's | 0.0001<br>(****) | 2C |
|  | 17-19 | 98 ± 6 | 102 (54) | Y |  |  |  |
|  | 19-21 | 164 ± 10 | 150 (58) | N |  |  |  |
|  | 25°C | 85 ± 3 | 84 (31) | Y |  |  |  |
| *Canton-S*<br>(gabazine<br>control) | 15-17 | 85 ± 5 | 88 (48) | Y | Kruskal-Wallis,<br>Dunn's | 0.0001<br>(****) | 2D |
|  | 17-19 | 153 ± 12 | 134 (119) | N |  |  |  |
|  | 19-21 | 91 ± 9 | 88 (48) | Y |  |  |  |
|  | 25°C | 81 ± 5 | 78 (60) | N |  |  |  |
| *Pan-neuronal-*<br>*Gal4 (Auxin-*<br>*Gal80)>UAS-*<br>*Rdl* | 15-17 | 182 ± 8 | 178 (90) | Y | One-way<br>ANOVA,<br>Tukey's | 0.0001<br>(****) | 3A |
|  | 17-19 | 82 ± 4 | 82 (44) | Y |  |  |  |
|  | 19-21 | 100 ± 5 | 97 (48) | Y |  |  |  |
|  | 25°C | 89 ± 4 | 90 (68) | Y |  |  |  |
| *Pan-neuronal-*<br>*Gal4 (Auxin-*<br>*Gal80)>UAS-*<br>*Gad1 RNAi* | 15-17 | 120 ± 5 | 108 (34) | N | Kruskal-Wallis,<br>Dunn's | 0.0001<br>(****) | 3B |
|  | 17-19 | 107 ± 4 | 105 (38) | N |  |  |  |
|  | 19-21 | 184 ± 9 | 189 (99) | Y |  |  |  |
|  | 25°C | 76 ± 2 | 72 (31) | Y |  |  |  |
| *muscle-GFP,*<br>*OK371*<br>(control,<br>diazepam,<br>gabazine) | - | - | - | Y | Mixed<br>effects<br>analysis,<br>Tukey's | 0.095<br>(ns) | 4B |

## Results

### A CP exists between 17-19 h AEL

The *Drosophila* embryonic CP represents a window of heightened plasticity during which induced activity-perturbation (e.g., mediated by optogenetics or drug exposure) produces a permanent larval seizure phenotype that is characterised by a greater duration of seizure-like behaviour in response to a strong electrical stimulus (Giachello and Baines, 2015, Hunter et al., 2024). To provide a more realistic mode of activity-perturbation, one that a developing embryo might experience in the wild, we used acute temperature stress (Doran et al., 2025). Initially, we sought to replicate previous work and validate the timing of the CP, when embryos are exposed to a temperature of 32°C. Embryos of successive developmental ages (15, 17 or 19 h AEL, respectively) were heated to 32°C for 2 h, and thereafter reared at 25°C until the wall-climbing L3 stage. Larve were then individually subjected to electroshock-induced seizure. Controls included non-manipulated embryos collected simultaneously and reared at 25°C throughout development (negative control) and *para*^*Bss*^ seizure mutants reared at 25°C throughout development (positive control). We observed a significant difference between seizure recovery times (RTs) for embryos that had been heat-stressed at different developmental ages (one-way Kruskal-Wallis ANOVA: *H*_(4, N=48)_ = 28.59, *p* < 0.0001). Dunn’s *post hoc* analysis identified the greatest effect at 17-19 h AEL compared to 15-17 (*p* < 0.0001) and 19-21 h AEL (*p* < 0.0001). A difference between 15-17 h AEL and 19-21 h AEL (*p* > 0.99) failed to reach significance, relative to non-manipulated controls (Figure 1). These data show that a 32°C heat stress, when applied during the canonical CP window of 17-19 h AEL, but not 2-h before or after, is sufficient to cause a strong seizure phenotype when tested several days post-manipulation at the end of the larval stage, indicative of a lasting reduction in network stability. This validates previous findings that identified an embryonic CP of nervous system development from 17-19 h of embryonic development (Giachello and Baines, 2015).

**Figure 1.**
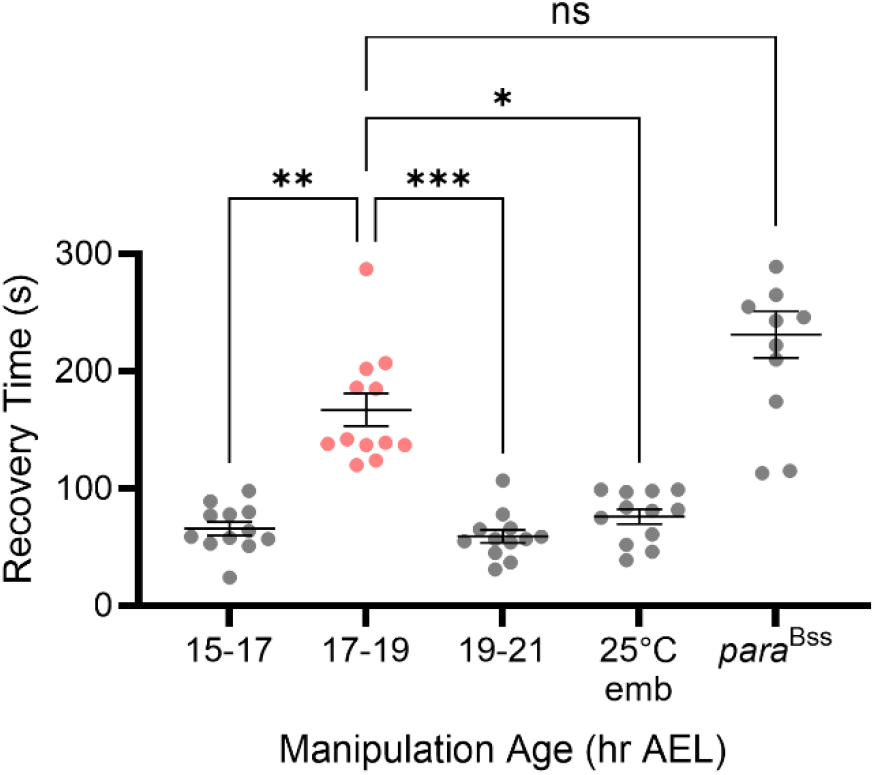
A critical period occurs at 17-19 h AEL. Seizure recovery times (RTs) of L3 larvae subjected to electroshock, derived from embryos exposed to 32°C at different developmental ages, and appropriate controls. Dunn’s *post hoc* analyses revealed a significant increase in RT for larvae derived from embryos exposed to 32oC heat stress during 17-19 h AEL compared to 15-17 or 19-21 h AEL (N = 12, respectively). RTs for wildtype larvae transiently exposed to 32oC between 15-17 or 19-21 h AEL were not significantly different compared to larvae derived from wildtype embryos maintained at 25°C throughout. Controls consisted of embryos not exposed to heat stress (25*°*C emb) or *para*^*Bss*^ seizure mutants (positive controls) maintained at 25°C throughout. \*\**p* < 0.01, \*\*\**p* < 0.001, \*\*\*\**p* < 0.0001. Data are presented as mean ± SEM.

### Pharmacological manipulation of GABAergic signalling alters the timing of CP opening

Increased signalling of intracortical GABA is considered necessary for the opening of CPs during mammalian neural development, most notably that of ocular dominance in the developing visual cortex (Fagiolini and Hensch, 2000). The initial expression of GAD1, hence the synthesis of GABA, in *Drosophila* (∼16-17 h AEL) immediately precedes the onset of the *Drosophila* embryonic locomotor CP (Kuppers et al., 2003). Therefore, we investigated whether onset of GABAergic signalling regulates the opening of the embryonic locomotor CP.

To potentiate GABAergic signalling, we exposed developing embryos to the GABA agonist diazepam. This was achieved by feeding gravid female flies with diazepam-supplemented (0.5 mM) food for 3 days. Subsequently, diazepam-exposed embryos were separated into three groups, and aged to expose them to a transient 32°C heat stress either from 15-17, 17-19 or 19-21 h AEL. Resultant wall-climbing L3 were subjected to electroshock. As expected, electroshock produced different RTs between the three groups and a control reared at 25°C throughout (one-way Kruskal-Wallis ANOVA: *H*_(4, N=96)_ = 19.14, *p* = 0.0003). Dunn’s *post hoc* analysis identified 15-17 h AEL (154 s ± 14) as the interval with the greatest RT, compared to 17-19 h (80 s ± 5; *p* = 0.0003), 19-21 h AEL (99 s ± 12; *p* = 0.007) or constant 25°C (97 s ± 8, *p* = 0.0102) (Figure 2A). Thus, in the presence of diazepam, the opening of the CP shifted earlier. The period duration, however, was not seemingly affected. To control for potential vehicle effects, separately aged collections of embryos were exposed to ethanol and heat-stressed in an identical manner. The result showed no effect with the biggest RT being seen at the expected 17-19 h AEL window (one-way Kruskal-Wallis: *H*_(4, N=106)_ = 35.40, *p* < 0.0001) (Figure 2B).

**Figure 2.**
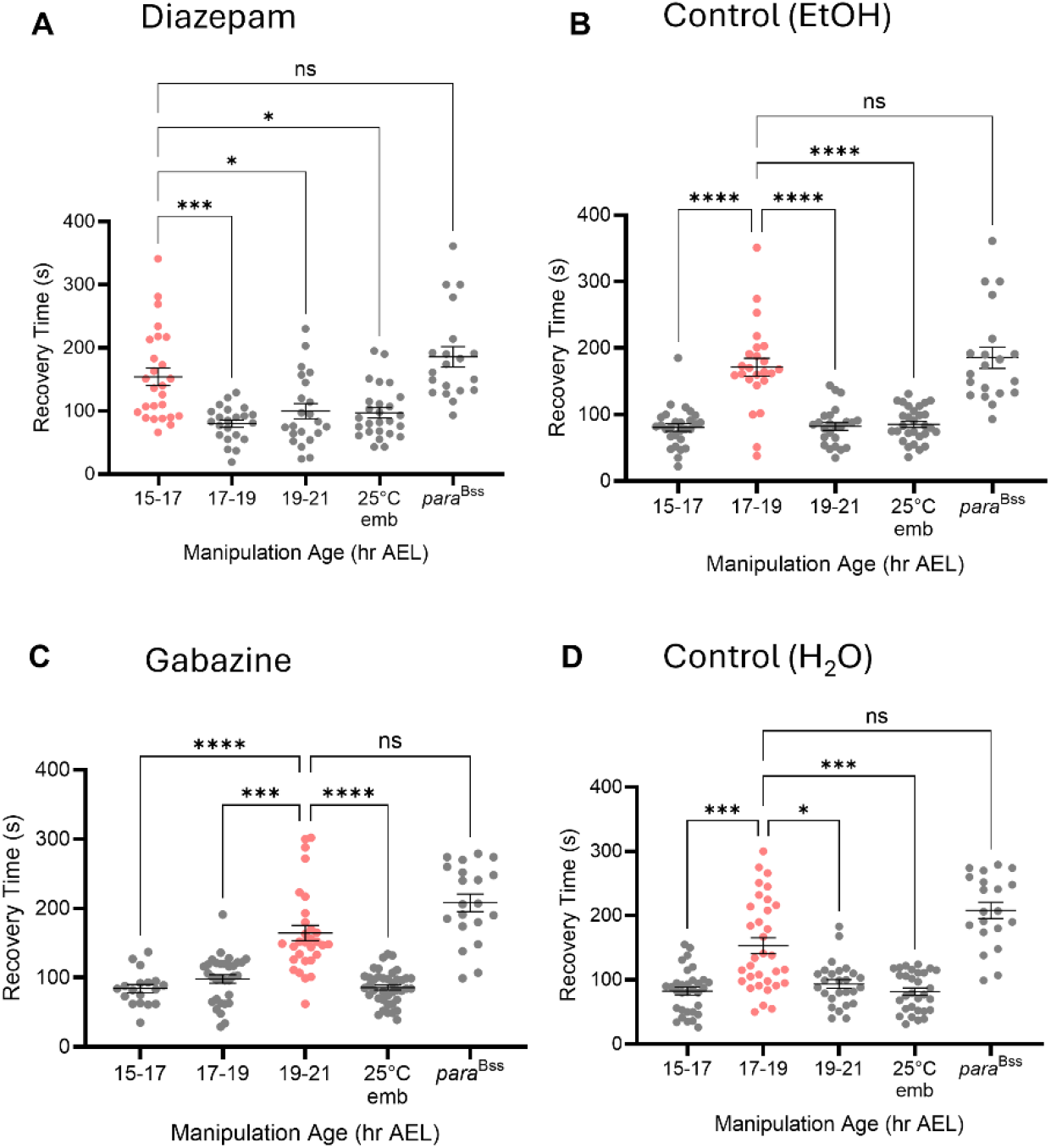
Pharmacological manipulation of GABA signalling alters the timing of CP opening. Across all presented experiments, drug or control solution was administered to embryos via transfer from gravid adult females. Embryos were then exposed to 32°C at either 15-17, 17-19 or 19-21 h AEL and then reared at 25°C and electroshocked at L3. ***A***, CS embryos fed 0.5 mM diazepam (GABA^A^ agonist). Larvae derived from embryos heat-stressed at 15-17 h AEL (N = 26) exhibited the greatest RT, whereas RTs for those manipulated at 17-19 (N = 22) or 19-21 h (N = 22) were comparable to the 25°C control (N = 26). ***B***, CP timing validation with ethanol-fed CS, controlling for potential vehicle-induced effects. Larvae derived from embryos heat-stressed at 17-19 h AEL (N = 24) exhibited significantly increased RTs compared to 15-17 (N = 24), 19-21 h (N = 28) and the 25°C control (N = 30). ***C***, CS embryos fed 0.5 mM gabazine (GABA^A^ antagonist). Larvae derived from embryos heat-stressed at 19-21 h AEL (N = 30) exhibited RTs significantly greater than those exposed at 17-19 (N = 32), 19-21 h (N = 30), or the 25°C control (N = 40). ***D***, CP timing validated with water-fed CS, controlling for potential vehicle-induced effects with gabazine. Larvae derived from embryos exposed to 32°C at 17-19 h AEL (N = 33) exhibited significantly increased RTs. Statistical differences were identified by a series of one-way Kruskal-Wallis ANOVA followed by Dunn’s *post hoc* analyses. \**p* < 0.05, \*\**p* < 0.01, \*\*\**p* < 0.001, \*\*\*\**p* < 0.0001. All data are presented as mean ± SEM.

To further validate the role of GABAergic signalling in CP opening, 0.5 mM gabazine, a competitive antagonist of the *Drosophila* GABA_A_ receptor (Hosie and Sattelle, 1996), was administered to gravid females and separately aged embryos were simultaneously heat-stressed as described above. As expected, administering a 32°C heat stress to groups of embryos from gabazine-fed gravid females, at the respective developmental windows, produced different seizure RTs following electroshock (one-way Kruskal-Wallis: *H*_(4, N=120)_ = 49.63, *p* < 0.0001). Dunn’s *post hoc* analysis revealed a marked increase in RT in larvae derived from embryos heat-stressed at 19-21 h AEL (164 s ± 10), i.e. two hours later than the normal embryonic CP; as compared with those having received 32°C heat stress at 15-17 h (84 s ± 5; *p* < 0.0001), 17-19 h (98 s ± 6; *p* < 0.0001) or 25°C controls (85 s ± 3, *p* < 0.0001) (Figure 2C). Controls, using a water vehicle, showed the expected CP window at 17-19 h AEL (Figure 2D). Thus, antagonism of GABA produced the opposite effect to diazepam, in that it delayed the opening of the embryonic CP.

### Genetic manipulation of embryonic GABAergic signalling alters the timing of CP opening

To validate the results observed using pharmacological manipulation of GABAergic signalling, we utilised a complementary, but independent, genetic approach. To mimic the activity of diazepam, we pan-neuronally over-expressed *resistance to dieldrin (rdl*) *that encodes a* GABA_A_ receptor subunit. This subunit is able to form functional homo-oligomeric receptors (Buckingham et al., 1996, Liu et al., 2007). To avoid potential masking effects of Gal4 expression during later larval development, we utilised the auxin-inducible gene expression system (AGES) to restrict transgene expression to embryonic and early larval development (McClure et al., 2022). As anticipated, the L3 electroshock assay produced different RTs across the three heat-stressed groups and 25°C controls (one-way ANOVA_*F*(3, 136)_ = 63; *p* < 0.0001). A Tukey’s *post hoc* test indicated embryos manipulated at 15-17 h AEL (182 s ± 8) exhibited a significant increase in RT relative to those manipulated at 17-19 h (82 s ± 4; *p* < 0.0001), 19-21 h (100 s ± 5; *p* < 0.0001) or 25°C controls (89 s ± 4, *p* < 0.0001) (Figure 3A). These data show that genetic potentiation of GABAergic signalling, via *rdl* overexpression, confers a premature opening of the CP. This mimics the effects observed following diazepam exposure.

**Figure 3.**
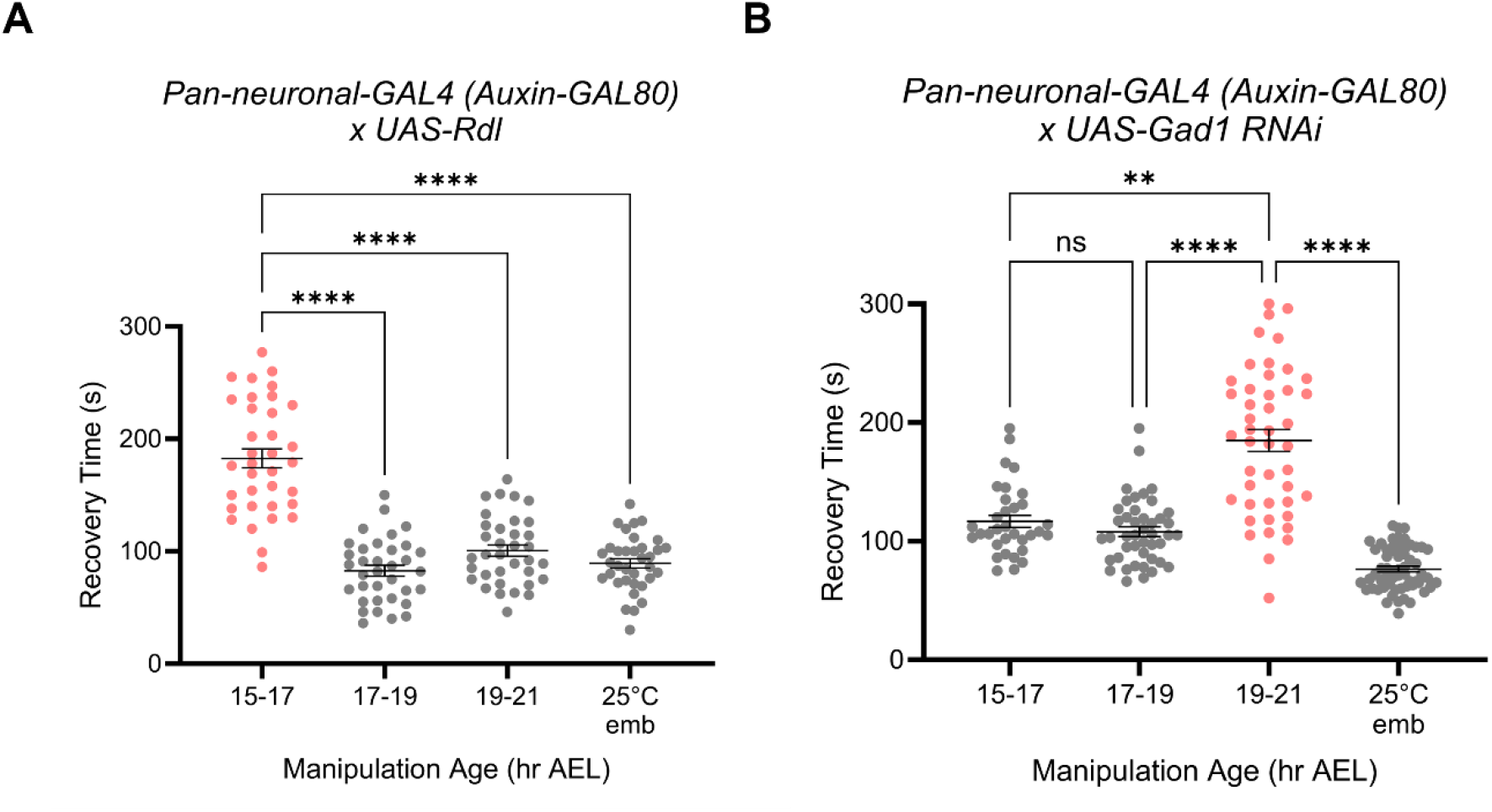
Genetic manipulation of GABA signalling alters CP opening. Across all experiments embryos were exposed to 32°C at either 15-17, 17-19 or 19-21 h AEL (respectively) and then reared at 25°C and electroshocked at L3. Controls were maintained at 25°C throughout (25°C emb). ***A***, Pan-neuronal expression of the GABA^A^ receptor (*nSyb-Gal4 (Auxin-GAL80)>UAS-rdl*). RTs, as indicated by a one-way ANOVA and Tukey’s *post hoc* test, were significantly greater for larvae derived from embryos heat-stressed at 15-17 h AEL (N = 35, respectively). ***B***, Pan-neuronal knockdown of *Gad1* (*nSyb-Gal4 (Auxin-GAL4)>UAS-Gad1 RNAi*). RTs, as indicated by a one-way Kruskal-Wallis ANOVA and Dunn’s *post hoc* test, were significantly greater in larvae derived from embryos heat-stressed at 19-21 h AEL (N = 45), compared to larvae heated at 15-17 h (N = 33), 17-19 h (N = 44) or 25°C controls (N = 60). \*\**p* < 0.01, \*\*\*\**p* < 0.0001. All data are presented as mean ± SEM.

To delay onset of GABAergic signalling, we pan-neuronally knocked down the expression of *Gad1* – the *Drosophila* GABA-synthetic enzyme – throughout embryogenesis. Embryos were collected, aged and transiently heat-stressed at 32°C during the developmental windows of interest, as described previously. The selective knockdown of *Gad1* produced different RTs in L3 manipulated across the three developmental intervals and 25°C controls (one-way Kruskal-Wallis: *H*_(4, N=182)_ = 104.5, *p* < 0.0001). Dunn’s *post hoc* analyses show that RT was significantly greater in larvae derived from embryos heat-stressed at 19-21 h AEL (184 s ± 9), compared to those exposed at 15-17 h (120 s ± 5 *p* = 0.002), 17-19 h (107 s ± 4 *p* < 0.0001) or 25°C controls (76 s ± 2, *p* < 0.0001) (Figure 3B). These data suggest that CP opening is delayed when GABA synthesis is reduced and thus GABAergic signalling downregulated, as in the mammalian CP for ocular dominance. Taken together, our data provides convincing evidence that the mechanism by which CPs open is conserved between mammals and *Drosophila*.

### Pharmacological manipulation of GABA signalling does not alter progression of patterned muscle activity during the CP

Recent studies suggest that patterned activity of body-wall muscles initiates neural development and possibly contributes to the timing of the embryonic locomotor CP (Carreira-Rosario A et al., 2021, Zeng X et al., 2021). We therefore questioned whether altered CP timing that results from manipulation of GABAergic signalling might be caused by altered timings of body-wall muscle activity. To determine this, we imaged embryos expressing GFP at their muscle z-discs (these occur between segmental muscles and provide a convenient measure of muscular contraction). We quantified the progression of coordinated locomotor activity (Figure 4A), beginning at 16 h AEL (i.e., before the CP opens) and extending to 19 h AEL (when the CP closes). In addition to control embryos, we also analysed embryos from gravid female flies that were fed drug-supplemented (0.5 mM) food for 3 days to potentiate (diazepam) or attenuate (gabazine) GABAergic signalling, respectively. A mixed-effects analysis showed no significant effects of treatment on the number of contraction waves exhibited at any developmental age tested (16, 17, 18 and 19 h AEL) (*F*_(2, 27)_ = 2.565, *p* = 0.1) (Figure 4B). Therefore, exposure to either diazepam or gabazine did not alter the timing of patterned muscle activity. These data imply that manipulation of GABAergic signalling, sufficient to alter CP timing, does not alter the developmental timeline of body-wall muscle contractions.

**Figure 4.**
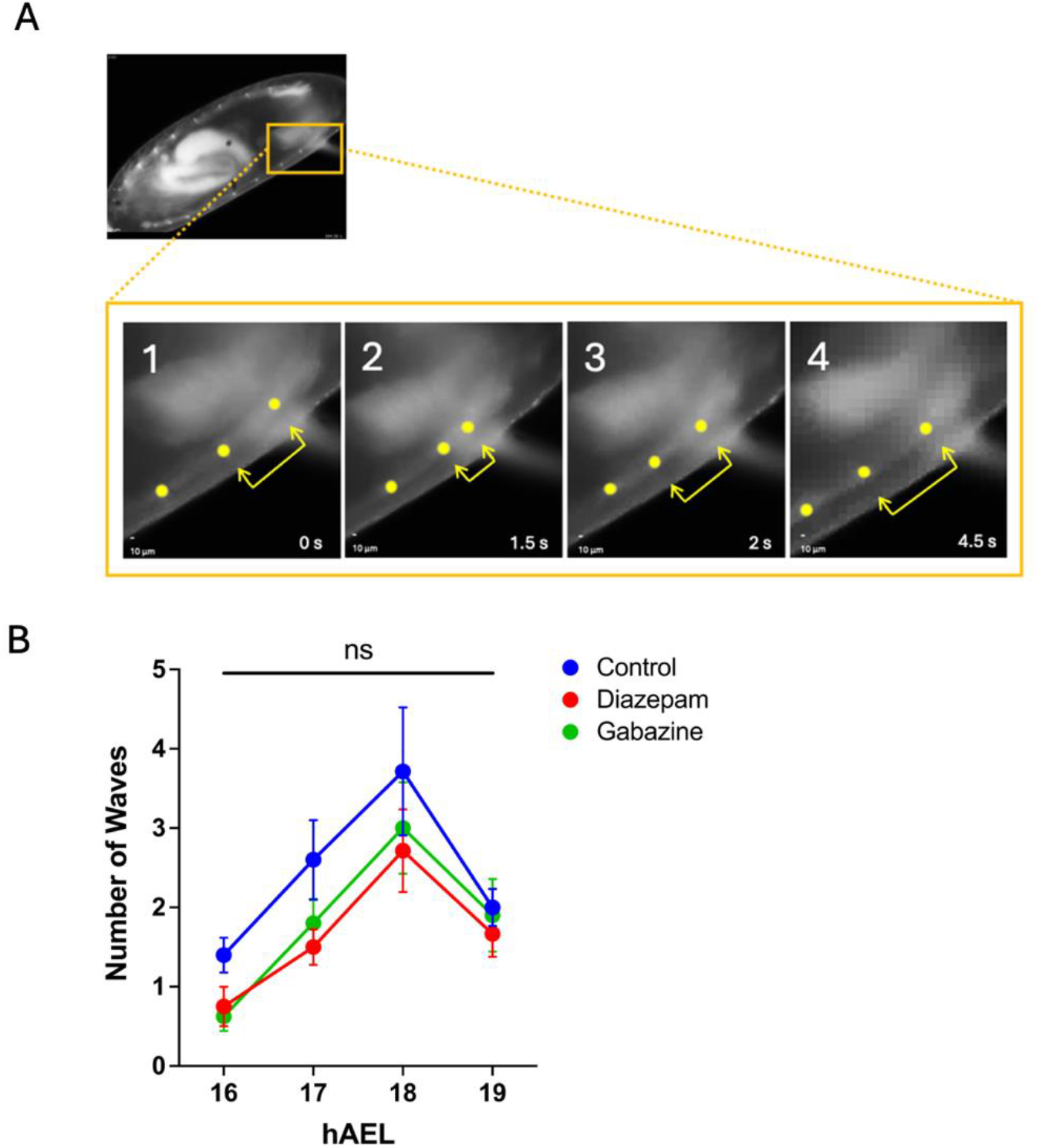
Pharmacological manipulation of GABA signalling does not alter onset or progression of patterned muscle activity. Drugs were administered to embryos via transfer from gravid adult females. Staged 13 h AEL embryos carrying muscle-GFP targeting z-discs were selected and imaged between 16-19 h AEL. ***A***, Images show an example of muscle z-discs contracting and relaxing. The sequence of these events (frames 1-4) throughout the whole embryo length were considered a coordinated ‘wave’. To increase contrast, yellow dots have been overlaid on z-discs, and yellow arrows illustrate the increasing and decreasing distance between two adjacent z-discs. ***B***, Patterned muscle activity was quantified as the number of waves exhibited by embryos during a 3 min recording, at every hour between 16 and 19 h AEL. No significant differences were found between embryos fed diazepam or gabazine and controls (mixed-effects analysis, *p* = 0.1, N = 7-10 per developmental age and per treatment group). All data are presented as mean ± SEM.

## Discussion

The contribution of CPs to neural circuit development remains to be fully understood. An intriguing hypothesis is that, during such periods, developing networks measure, and where possible, compensate for unexpected influences that might otherwise alter the functionality of a mature circuit (Hunter et al., 2024). Indeed, experimental manipulation of activity during a variety of CPs, ranging from invertebrates to mammals, all consistently show permanent, and often detrimental, effects to mature circuit function (Coulson et al., 2022, Giachello and Baines, 2015, Hensch, 2005). To date, much of our understanding of the contribution of CPs to neural circuit development has been gained from studies in mammals, particularly the mammalian visual cortex (Hensch, 2005, Hensch and Quinlan, 2018). Whilst this elegant work has provided much insight, the complexity of mammalian sensory systems presents specific challenges. Thus, it is significant that CPs are found widely across phyla and, particularly, in the experimentally tractable fruit fly, *Drosophila*. The many advantages that accompany this model insect, from advanced genetics to a full larval connectome, offer the opportunity to fully explore the events that occur during a CP.

In this study, we show that the timing of CP opening in the development of the *Drosophila* locomotor circuitry is influenced by GABAergic signalling. There are several receptor types that bind GABA and mediate ionotropic (GABA_A_ and GABA_C_) or metabotropic (GABA_B_) transmission (Hosie et al., 1997). In *Drosophila*, the primary GABA_A_ receptor subunit, which can form homo-oligomeric receptors, is encoded by *rdl* (Buckingham et al., 1996, Liu et al., 2007). In our experiments, embryonic diazepam administration or overexpression of *rdl* was sufficient to prematurely open the locomotor CP (at 15-17 h AEL, instead of 17-19 h AEL). By contrast, embryonic administration of gabazine, a competitive antagonist of the of GABA_A_ receptor, or RNAi-mediated knockdown of *Gad1*, delayed CP opening to 19-21 h AEL. Our observations suggest that in *Drosophila*, as in mammals, increased GABAergic signalling is sufficient to open a CP (Fagiolini and Hensch, 2000). Thus, GABAergic signalling appears to be a key regulator of CP timing across phyla.

To investigate the recent proposal that functional embryonic nervous system development in *Drosophila* is initiated by a change from spontaneous to patterned activity of body wall muscles (Carreira-Rosario A et al., 2021, Zeng X et al., 2021), we tracked the emergence of coordinated movement in embryonic muscles and how this is influenced by exposure to the GABA-altering compounds used in this study. We could not, however, observe significant change in the sequential appearance of coordinated segmental contractions that herald the onset of patterned activity. Thus, this suggests that CP opening is mediated by maturation of GABAergic signalling within the CNS, rather than changes in the timing of muscle contractions. GABA occupies a multifaceted niche during mammalian development. Indeed, early GABAergic signalling is depolarising and switches to hyperpolarising in most neurons during development (Peerboom and Wierenga, 2021). This switch in valence is mediated by the preferential downregulation of the Na-K-Cl cotransporter 1 (NKKC1; Cl^-^ importer), and the upregulation of the K-Cl cotransporter 2 (KCC2; Cl^-^ exporter). Initially, however, when glutamatergic signalling is minimal, depolarising GABA directs the sequential transition between several key phases of neurogenesis. The earliest occurrence of GABAergic activity, in developing mouse, is autocrine signalling via GABA_A_ receptors expressed by embryonic neural crest stem cells (Andang et al., 2008, Wang and Kriegstein, 2008). Subsequently, GABA promotes the proliferation of primary neural stem cells and prevents the proliferation of neural progenitor cells to ensure an appropriate balance between proliferation and differentiation as the brain develops (Young et al., 2012, Haydar et al., 2000). Depolarising GABAergic signalling also regulates the migration of newborn neurons to appropriate neuroanatomical regions (Luhmann et al., 2015), promotes Ca^2+^-mediated neurite outgrowth (Cancedda et al., 2007, Sernagor et al., 2010), and regulates synapse generation and maturation (Oh et al., 2016). Therefore, given the extensive actions of GABA and the fundamental role of CPs during neural development, it is perhaps unsurprising that this neurotransmitter regulates CP timing.

In *Drosophila*, the expression of GABA follows that of *Gad1* expression, which first becomes identifiable at ∼16 h AEL (Kuppers et al., 2003). This timing coincides with the first appearance of synaptic currents in motoneurons of the developing locomotor circuitry (Baines and Bate, 1998). Thus, in many respects the opening of the locomotor CP occurs relatively early in development, at a time when the locomotor network is only just beginning to show first signs of functionality. Our previous work has suggested that this CP, when open, integrates activity present in the CNS, summing both excitatory and inhibitory signalling, to properly encode the excitation:inhibition balance of the mature larval locomotor circuitry. Activity-manipulation during this period is sufficient to permanently alter this balance, and when extreme, to leave the network vulnerable to induced seizures (Hunter et al., 2024).

In summary, we show that CP opening, in a *Drosophila* circuit, is timed by the maturation of GABAergic signalling. This highlights the shared role of GABA in CP opening between *Drosophila* and mammals, strengthening the tractability of *Drosophila* to fully explore the contribution of a CP to neural circuit development.

## Acknowledgements

This work was supported by funding from a Joint Wellcome Trust investigator award to R.A.B. and M.L. (Grant 217099/Z/19/Z). Work on this project benefited from the Manchester Fly Facility, established through funds from the University and the Wellcome Trust (Grant 087742/Z/08/Z).

## Data and resource availability

All relevant data and details of resources can be found within the article.

## Author Contributions

Conceptualization, J.C and R.A.B.; Methodology, J.C and M.H.R and M.L.; Investigation, J.C and M.H.R.; Writing – Original Draft, J.C and M.H.R.; Writing – Review & Editing, J.C, M.H.R. M.L and R.A.B.; Funding Acquisition, R.A.B. and M.L; Resources, R.A.B. and M.L.; Supervision, R.A.B.

